# The Salivary Microbiome: Analysis of by Pyrosequencing and the Relationship with *Helicobacter pylori* Infection

**DOI:** 10.1101/505115

**Authors:** Yingjie Ji, Xiao Liang, Hong Lu

## Abstract

**Backgrounds:** There have been reports of *Helicobacter pylori* (*H. pylori*) in the oral cavity and it has been suggested that the oral cavity may be a reservoir for *H. pylori* reflux from the stomach.

**Objectives:** High-throughput pyrosequencing was used to assess the structure and composition of oral microbiota communities in individuals with or without confirmed *H. pylori* infection.

**Methods:** Saliva samples were obtained from 34 *H. pylori* infected and 24 *H. pylori* uninfected subjects. Bacterial genomic DNA was extracted and examined by pyrosequencing by amplification of the 16S rDNA V3-V4 hypervariable regions followed by bioinformatics analysis. Saliva sampling was repeated from 22 of the 34 *H. pylori* infected subjects 2 months after *H. pylori* eradication.

**Results:** High-quality sequences (2,812,659) clustered into 95,812 operational taxonomic units (OTUs; 97% identity), representing 440 independent species belonging to 138 genera, 68 families, 36 orders, 21 classes, and 11 phyla. Species richness (alpha diversity) of *H. pylori* infected subjects was similar to that of uninfected subjects. Eradication treatment decreased saliva bacterial diversity. Beta diversity analysis showed that the salivary microbial community structure differed between *H. pylori* infected and uninfected subjects both before and after *H. pylori* eradication.

**Conclusions:** Salivary microbiota diversity was similar in *H. pylori* infected and uninfected individuals. Antibiotic therapy was associated with a decline in salivary bacterial diversity. Both *H. pylori* infection and its eradication caused the oral microbiota alterations in community and structure. The present of *H. pylori* in oral cavity was not related with its infection status in stomach.

**Trial registration:** ClinicalTrials.gov, Identifier: NCT03730766

**Importance:** The oral cavity plays a vital role in *Helicobacter pylori* transmission among human. High-throughput pyrosequencing of the 16S rDNA V3-V4 hypervariable regions was used to assess the structure and composition of oral microbiota communities in individuals with or without confirmed *Helicobacter pylori* infection. We show that both *Helicobacter pylori* infection and eradication cause microbiota alterations in the oral microbiota. Prior studies report detection of *Helicobacter pylori* in the oral cavity by polymerase chain reaction. We show that the presence of *Helicobacter pylori* in the oral cavity is unrelated with its infection status in the stomach.

## Introduction

*Helicobacter pylori* (*H. pylori*) is a Gram-negative bacterium that colonizes the human gastric epithelium. It belonged to *Helicobacter* genus, *Helicobacteraceae* family, *Campylobacterales* order, *Epsilonproteobacteria* class, and *Proteobacteria* phyla. *H. pylori* infection is characterized by mucosal inflammation (gastritis) and may result in peptic ulcer disease or gastric adenocarcinoma (1). *H. pylori* is transmitted between humans by a variety of routes including gastro-oral and fecal-oral mechanisms that include contaminated water and food. It has also been postulated that oral cavity may play a role in *H. pylori* transmission and possibly act as a reservoir (2). For example, *H. pylori* has been detected in the oral cavity using the polymerase chain reaction (PCR). *H. pylori* has also been successfully cultured from saliva from individuals with positive results of both a saliva *H. pylori* antigen test and *H. pylori* flagella test (3).

The oral cavity is one of the most complex and largest microbial habitats that harbors hundreds of different bacteria which play important roles in maintaining oral homeostasis and influencing the development of both oral and systematic diseases (4). Many factors in the oral environment including intraoral pH and salivary iron concentration have been reported to have significant relationships with oral microbial communities (5). However, there was few reports about *H. pylori* and its relationship with the microbial community structure in human saliva.

Currently, most oral bacteria species cannot be cultivated *in vitro* using traditional cultivation methods requiring the use of molecular biological techniques, such as checkerboard hybridization, microarray chips, and the quantitative real-time PCR (6) to identify and classify the currently uncultivable bacteria. However, many low-abundance bacteria species still cannot be detected using these approaches which impedes the comprehensive and in-depth understanding of oral bacteria diversity. In this study, we used amplicon pyrosequencing of 16S rDNA V3-V4 hypervariable regions to define the bacterial composition, abundance, and structure of salivary microbiome in people with and without active *H. pylori* infections. In addition, we also characterized the salivary biodiversity of a subgroup of subjects before and after the *H. pylori* eradication.

## Methods

### Subjects and Sample Collections

This study was performed in accordance with the recommendations of the Ethics Committee of the Renji Hospital of Shanghai Jiao Tong University. All subjects gave written informed consent in accordance with Declaration of Helsinki.

The samples were collected in Renji Hospital of Shanghai Jiao Tong University, China, from August to November in 2018. A total of 58 subjects were recruited, including 34 subjects with *H. pylori* infection and 24 uninfected subjects. We first conducted a cross-sectional study of the salivary microbiota of 34 *H. pylori* infected and 24 uninfected subjects. A prospective study was then performed in a subgroup of 22 subjects with *H. pylori* infection who underwent salivary analysis both before and after successful *H. pylori* eradication.

All subjects received both endoscopy and ^13^C urea breath test (^13^C-UBT) before enrollment. The *H. pylori* infection status was confirmed by positive rapid urease test (RUT), histology and ^13^C-UBT. Absence of infection was defined as negative results for all tests (i.e., RUT, histology and ^13^C-UBT). *H. pylori* infected subjects received eradiation therapy consisting of esomeprazole 20 mg b.i.d., bismuth potassium citrate 600 mg b.i.d, amoxicillin 1000 mg t.i.d., and metronidazole 400 mg t.i.d. for 14 days. *H. pylori* eradication was confirmed by ^13^C-UBT at least 6 weeks after the end of treatment. Saliva samples were collected from 22 subjects both before and 2 months following successful *H. pylori* eradication. Subjects were characterized into four groups. *H. pylori* uninfected group (uninfected) (n=24), *H. pylori* infected group (infected) (n=34), and Pre-eradicated *H. pylori* infected group (pre-eradicated) (n=22) and a successful eradicated group (eradicated) (n=22). Inclusion criteria of subjects were: age of 20-65 years old male or female, with good oral hygiene (including brushing teeth twice a day) and with no bad eating habits (7). Exclusion criteria included: 1) the presence of dental carious or any untreated cavitated carious lesions and oral abscesses, 2) periodontal disease or periodontal pockets ≥4 mm, 3) the use of antibiotics or PPI within 2 months before the study, 4) previous diagnosis of a serious systemic diseases (such as diabetes, hypertension or cardiopathy) or any diseases affecting oral health (such as Sjogren’s syndrome or any disease characterized by xerostomia), 5) pregnancy of breastfeeding, and 6) smoking or alcohol drinking. The detailed clinical parameters of the 58 subjects are shown in Table S1.

### Salivary sampling

Sampling was performed in the morning 2 hours after eating. Saliva samples were collected from each subject according to the Manual of Procedures for Human Microbiome Project (http://hmpdacc.org/resources/tools_protocols.php), with minor modifications. Approximately 3-4 ml of non-stimulated saliva was collected in two sterile, labeled 2 mL Eppendorf tubes, which were immediately placed on ice. Within 3 hours of collection, samples were transported on ice and stored at −80°C until use (8).

### DNA Extraction and Pyrosequencing

DNA was extracted from the saliva samples using the E.Z.N.A. ^®^ Soil DNA Kit (OMEGA, USA), following the manufacturer’s instructions, and stored at −20°C prior to further analysis. PCR amplification of the bacterial 16S rDNA hypervariable V3-V4 region was performed using the forward primer 338F (5’-ACTCCTACGGGAGGCAGCA-3’) and the reverse primer 806R (5’-GGACTACHVGGGTWTCTAAT-3’). Sample-specific 7-bp barcodes were incorporated into the primers for multiplex sequencing. Details of the barcodes are shown in Table S2. PCR amplification were performed on an ABI 2720 instrument (ABI, USA) with an initial denaturation at 98°C for 2 minutes, followed by 25 cycles of denaturation (15s at 98°C), annealing (30s at 55°C), extension (30s at 72°C), and ended with a final extension (5 min at 72°C). PCR amplicons were purified with Agencourt AMPure Beads (Beckman Coulter, Indianapolis, IN) and quantified using the Quant-iT PicoGreen dsDNA Assay Kit (Invitrogen, Carlsbad, CA, USA). Equimolar concentrations of purified amplicons were pooled in equal amounts. Subsequently, the paired-end 2×300 bp pyrosequencing was performed on the Illlumina MiSeq platform with MiSeq Reagent Kit v3 (Illlumina, USA), following the vendor’s standard protocols.

### Sequence Analysis

The Quantitative Insights Into Microbial Ecology (QIIME) pipeline was employed to process the sequencing data. Raw sequences were filtered to obtain high-quality sequences according to QIIME (9). The high-quality sequences were clustered into operational taxonomic units (OTUs) at 97% sequence identity by UCLUST (10). The representative sequences selected from each OTU were classified taxonomically by BLAST searching against the Human Oral Microbiome Database (HOMD), which provides a detailed record of the type, metabolism, and pathogenicity of oral bacteria (11). Then, an OTU table was further generated to record the OTU abundance of each sample and the taxonomic classification of these OTUs. Finally, to minimize the difference of sequencing depth across samples, the OTU table was modified by removing OTUs containing less than 0.001% of total sequences across all samples for further analysis (12).

### Bioinformatics and Statistical Analysis

Sequence data analyses were mainly performed using QIIME (version 1.8.0) and R packages (version 3.2.4). The alpha diversity analysis including Chao 1 richness estimator, Abundance-based Coverage Estimator (ACE) metric, Shannon diversity index, and Simpson index, were calculated at 97% identity using the QIIME (13). Ranked abundance curves were generated to compare both the richness and evenness of OTUs among samples. The beta diversity analysis including Nonmetric Multidimensional Scaling (NMDS), and unweighted UniFrac distances based principal coordinate analysis (PCoA), were performed using the R package to evaluate the similarity among various bacterial communities (14). The significance of differentiation of microbiota structure among groups was assessed by Adonis test (15). The taxonomy compositions and abundances were visualized by MEGAN (version 6.6.7) software (16). Linear discriminant analysis effect size (LEfSe) was used to compare the bacterial community structures between the samples from the patients with and without *H. pylori* infection, as well as before and after the eradication regimen, using the online Galaxy workflow framework (http://huttenhower.sph.harvard.edu/galaxy/) (17). Co-occurrence analysis among genera was performed among 50 most abundant genera. Correlations with |RHO| > 0.6 and P < 0.01 were visualized as co-occurrence network using Cytoscape (version 3.6.1) (18). Microbial functions were predicted by Phylogenetic investigation of communities by reconstruction of unobserved states (PICRUSt), based on high-quality sequences, and aligned to the Kyoto Encyclopedia of Genes and Genomes (KEGG) database (19).

### Data Access

All raw sequences were deposited in the NCBI Sequence Read Archive under accession number SRP167714.

## Results

### Global Sequencing Data

A total of 2,812,659 high-quality sequences (representing 79% of the total sequences) were acquired from the 80 saliva samples, with an average of 35,158 sequences per sample (ranging from 19,210 to 44,310). The average sequence length was 445 bp, with the maximum length being 548 bp and the shortest length being 136 bp (Figure S1). Clustering of all high-quality sequences at 97% identity resulted in 70,489 OTUs, which were BLAST-searched against the HOMD database for taxonomic assignments. After removing the low-credibility OTUs (together contributing only 6.7% of all sequences), a modified OTU table was obtained consisting of 95,812 OTUs with an average of 1,198 OTUs per sample (ranging from 697 to 1,584).

### Bacterial Abundance and Distribution

The bacterial distribution was characterized in terms of the relative taxonomic abundances. A total of 11 phyla, 21 classes, 36 orders, 68 families, 138 genera and 440 species were detected in the saliva samples. The taxonomic distributions of the predominant bacteria (relative abundance >1% of the total sequences) in subjects with and without *H. pylori* at different levels were shown in Figure 1. The 6 most abundant phyla were *Proteobacteria* (40.1% of the total sequences), *Firmicutes* (31.6%), *Bacteroidetes* (13.0%), *Actinobacteria* (7.4%), *Fusobacteria* (6.1%), and *TM7* (1.0%), together accounting for 99.2% of the total sequences. At genus level, saliva microbiota was dominated by *Neisseria, Streptococcus, Haemophilus, Veillonella*, and *Prevotella*, with average relative abundances of 20.2, 16.5, 10.5, 8.0, and 8.0%, respectively. The compositions in taxa of the microbial communities according to the tested sample groupings are provide in Figure S2.

**Figure 1:**
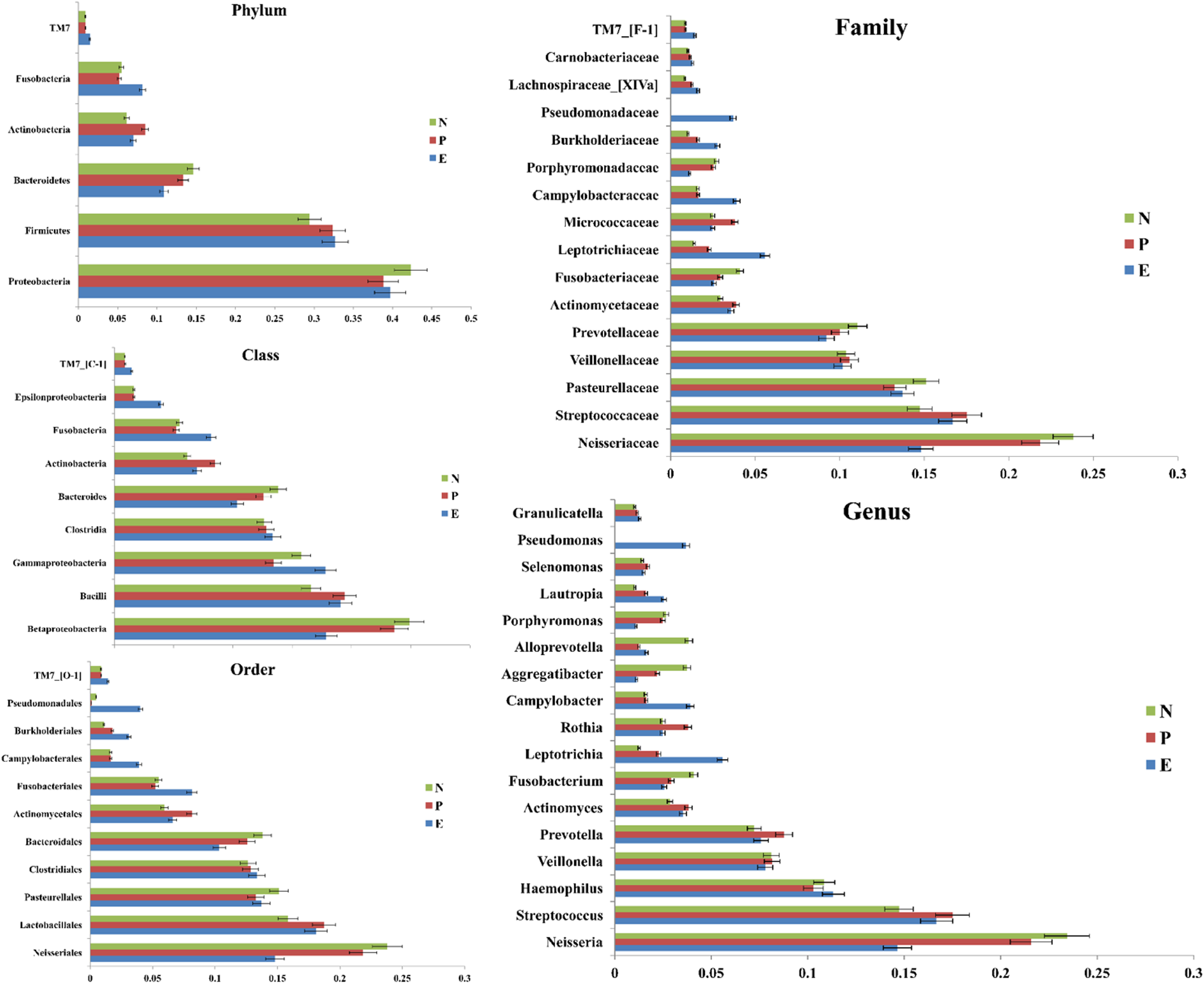
Distribution of the predominant bacteria at different taxonomic levels (phylum, class, order, family, and genus). The predominant taxa (>1% relative abundance) in each level are shown. N=uninfected group, P=infected group, E=eradicated group.

### Bacterial Diversity Analysis

The saliva microbiota richness, measured by numbers of observed OTUs, was similar in uninfected subjects and infected subjects (Figure S3A). The alpha diversity indices of Chao1, ACE, Shannon, Inverse Simpson, Good’s coverage, and Simpson’s evenness are shown in Table 1. The Shannon diversity index was higher in uninfected subjects than in infected subjects, but there was no significant difference between groups by *t*-test (1417.58 vs. 1393.60, p>0.05). Besides, the ACE richness index (1491.22 vs. 1465.97, p>0.05), Chao 1 richness estimator (1417.58 vs. 1393.60, p>0.05), and the Inverse Simpson diversity index (1.02 vs. 1.02, p>0.05) was also higher in uninfected subjects, with no significant difference, indicating the similar bacterial diversity of *H. pylori* uninfected saliva compared to the infected subjects. Good’s coverage estimator for each group was over 98%, indicating that the current sequencing depth was sufficient to saturate the bacterial diversity of saliva. In addition, Simpson’s evenness index indicated that the bacterial-community distribution in two groups was uneven, which was also observed in the rank-abundance curve (Figure S4).

**Table 1:** Alpha diversity indices for saliva bacteria in each group at 97% identity.

| Group | Chao1 |  | ACE |  | Shannon |  | Inverse Simpson |  | Coverage |  | Simpson even |  |
| --- | --- | --- | --- | --- | --- | --- | --- | --- | --- | --- | --- | --- |
|  | Mean | SE | Mean | SE | Mean | SE | Mean | SE | Mean | SE | Mean | SE |
| **N** | 1417.58 | 195.47 | 1491.22 | 210.41 | 7.94 | 0.36 | 1.02 | 0.01 | 0.98 | 0.01 | 0.05 | 0.02 |
| **P** | 1393.60 | 182.57 | 1465.97 | 196.06 | 7.89 | 0.42 | 1.02 | 0.01 | 0.98 | 0.00 | 0.05 | 0.01 |

SE, Standard Error. N=uninfected group, P=infected group.

### Bacterial Community Structures

To gain insights into the similarities in the bacterial community structures among uninfected and infected subjects, PCoA of beta diversity analysis was performed based on the unweighted UniFrac distances, which demonstrated different community structures among two groups (PERMANOVAR, pseudo-F: 1.49, p=0.033). As shown in Figure 2A, the overall microbial composition of infected subjects deviated from that of uninfected subjects. Furthermore, the results of NMDS based on the genus level classification exhibited clear segregations in community structures among groups (Figure S5A).

**Figure 2:**
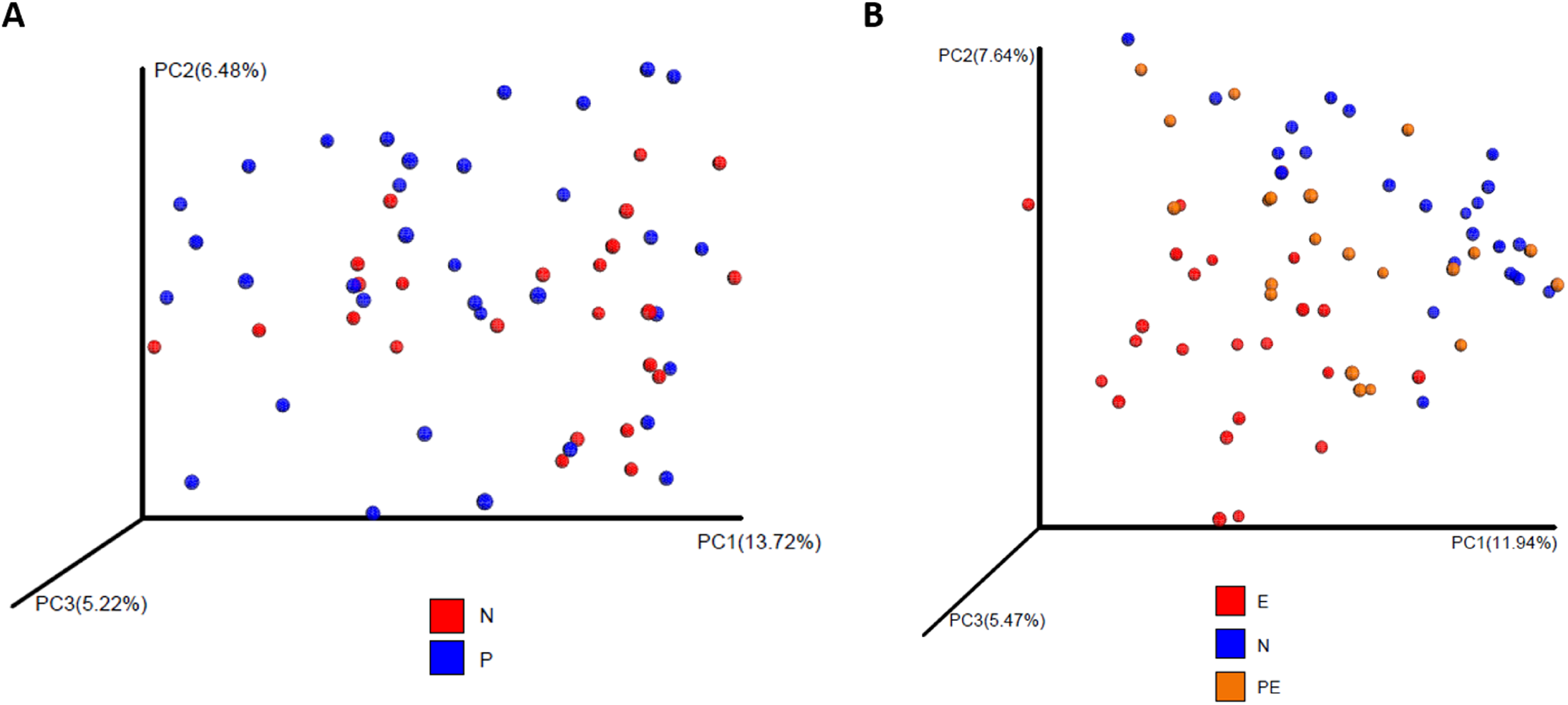
Principal coordinate analysis (PCoA) of unweighted UniFrac analysis. (A) PCoA analysis demonstrated that subjects of P group were significantly different from N group (PERMANOVAR, pseudo-F: 1.49, p=0.033). N=uninfected group, P=infected group. (B) PCoA analysis showed that the overall microbial composition showed significant difference between PE and E group (PERMANOVAR, pseudo-F: 3.34, p=0.001). E=eradicated group, N=uninfected group, PE=pre-eradicated group.

### Differential Microbiota Compositions

There were significant differences in the community compositions among two groups. As shown in Figure 3, a cladogram representation of significantly different taxa among groups was performed by LEfSe. The microbial composition was significantly different at the genus level, with 16 significantly different genera among the two groups. *Acinetobacter*, *Ralstonia, Leptothrix, Sphingomonas, Ochrobactrum, Afipia, Leptotrichia, Oribacterium*, and *Moryella* exhibited a relatively higher abundance in infected subjects, and can be considered *H*. pylori-enriched genera. *Alloprevotella, Aggregatibacter, Klebsiella, Leptotrichlaceae G_1_, Fusobacterium, Parvimonas*, and *Peptococcus* were relatively more abundant in uninfected subjects, which could be considered to be decreased in the infected group. These higher or lower expressed genera in infected subjects can be considered as *H. pylori*-associated genera (LAD >2, p< 0.05).

**Figure 3:**
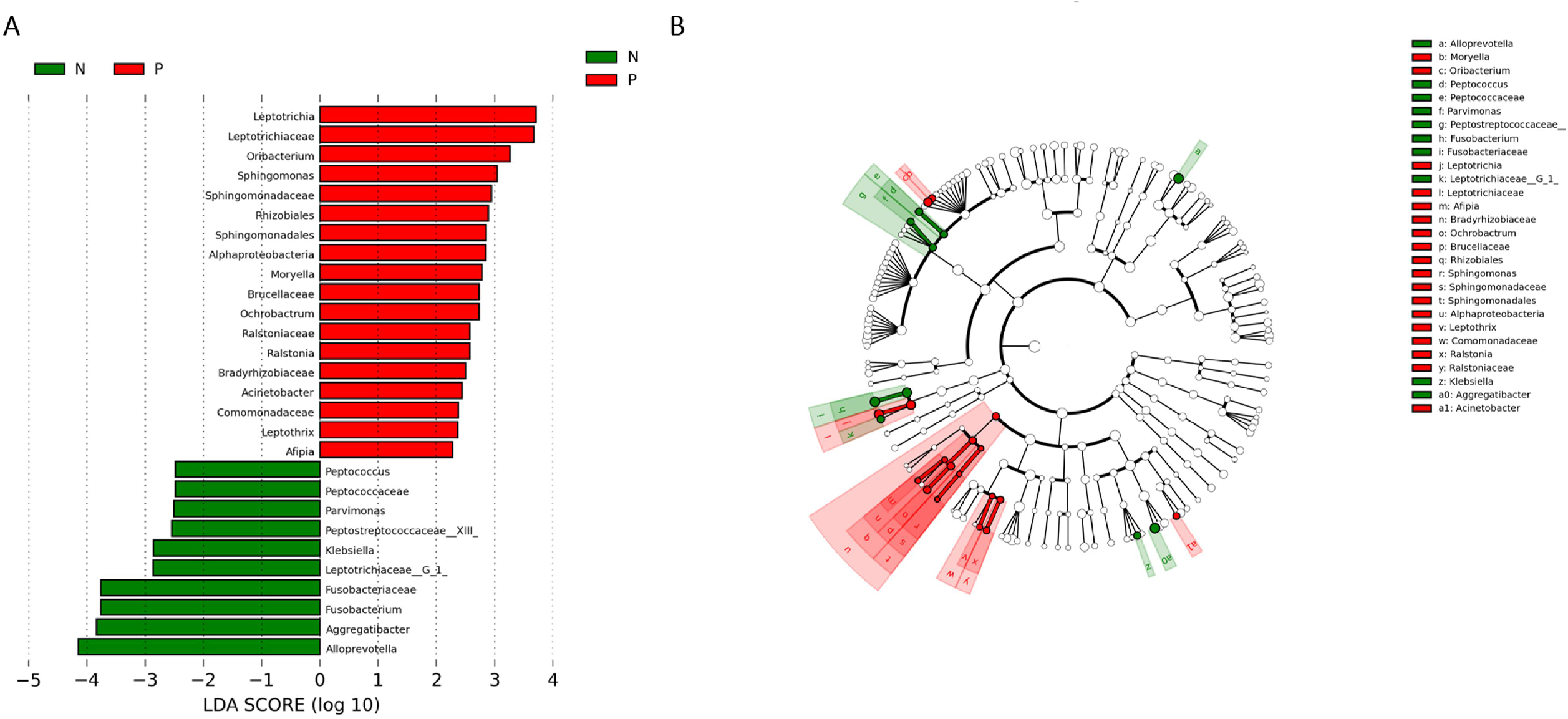
Comparison of microbial variations at the genus level, using the LEfSe online tool. (A) Histogram of the LDA scores for differentially abundant features among groups. The threshold on the logarithmic LDA score for discriminative features was set to 2.0. N=uninfected group, P=infected group. (B) Cladogram for taxonomic representation of significantly differences among groups. Differences are represented in the color of the most abundant taxa (red indicating P group, green indicating N group, and white indicating non-significant). N=uninfected group, P=infected group.

### Eradication therapy for *H. pylori* partially changed salivary microbiota

To determine the effects of *H. pylori* eradication therapy on saliva microbial composition, saliva samples from a subgroup of *H. pylori* infected subjects (n=22), were collected before (pre-eradicated group) and 2 months after treatment; saliva collected after successful eradication were classified into the eradicated group. The within-individual diversity in the samples from eradicated group was lower than pre-eradicated group (Figure S3B). The Shannon diversity index, ACE richness index, and Chao 1 richness estimator were higher in pre-eradicated subjects than in eradicated subjects, with a significant difference between groups by t-test (Shannon p=0.015, ACE p=0.003, Chao 1 p=0.002), indicating significant alteration in the within-individual diversity in samples from eradicated subjects compared to their baseline samples (Table 2).

**Table 2:** Alpha diversity indices for saliva bacteria in each group at 97% identity.

| Group | Chao1 |  | ACE |  | Shannon |  | Inverse Simpson |  | Coverage |  | Simpson even |  |
| --- | --- | --- | --- | --- | --- | --- | --- | --- | --- | --- | --- | --- |
|  | Mean | SE | Mean | SE | Mean | SE | Mean | SE | Mean | SE | Mean | SE |
| **PE** | 1350.56* | 200.94 | 1421.26* | 215.43 | 7.83* | 0.49 | 1.02 | 0.01 | 0.98 | 0.00 | 0.05 | 0.02 |
| **E** | 1143.95* | 215.08 | 1206.85* | 234.63 | 7.44* | 0.53 | 1.03 | 0.02 | 0.98 | 0.00 | 0.05 | 0.02 |

SE, Standard Error. Asterisk indicates significant difference ( $p < 0.05$ , Student's t-test). PE=pre-eradicated

| Group | Chao1 |  | ACE |  | Shannon |  | Inverse Simpson |  | Coverage |  | Simpson even |  |
| --- | --- | --- | --- | --- | --- | --- | --- | --- | --- | --- | --- | --- |
|  | Mean | SE | Mean | SE | Mean | SE | Mean | SE | Mean | SE | Mean | SE |
| **PE** | 1350.56<sup>*</sup> | 200.94 | 1421.26<sup>*</sup> | 215.43 | 7.83<sup>*</sup> | 0.49 | 1.02 | 0.01 | 0.98 | 0.00 | 0.05 | 0.02 |
| **E** | 1143.95<sup>*</sup> | 215.08 | 1206.85<sup>*</sup> | 234.63 | 7.44<sup>*</sup> | 0.53 | 1.03 | 0.02 | 0.98 | 0.00 | 0.05 | 0.02 |

SE, Standard Error. Asterisk indicates significant difference ( $p < 0.05$ , Student's t-test).
PE=pre-eradicated group, E=eradicated group.
\*Chao 1, \*ACE, and \*Shannon index between PE and E was statistically significant different ( $p < 0.05$ ).

The beta diversity using unweighted UniFrac showed significant differences in the overall microbial composition between pre-eradicated and eradicated groups (PERMANOVAR, pseudo-F: 3.34, p=0.001) (Figure 2B). The NMDS also exhibited clear segregations in community structures among groups (Figure S5B).

The relative difference of *H. pylori*-associated taxa was compared before and after eradication by LEFse analysis (Figure S6). Among the *H*. pylori-enriched genera, *Ralstonia, Leptotrichia, Sphingomonas*, *Leptothrix*, *Oribacterium*, and *Acinetobacter* increased after the eradication, while *Ochrobactrum* decreased after the successful eradication (p<0.05, paired Wilcoxon rank-sum test). Of the genera that decreased in infected subjects, *Alloprevotella, Aggregatibacter, Leptotrichlaceae G_1_, Parvimonas*, and *Fusobacterium* decreased after the eradication (p<0.05, paired Wilcoxon rank-sum test). Besides, we found that at phyla level, *Fusobacteria* increased after *H. pylori* eradication.

### *Helicobacter species* in the oral cavity

*H. pylori* was detected in 38 out of 80 saliva samples, occupying 0.0139% of all the total sequences (Figure 4). 12 of the 34 subjects in infected subjects (35.3%), 11 of 24 subjects in the uninfected group (45.8%), and 15 of 22 subjects in the eradicated group (68.2%) were found to possess *H. pylori* in the oral cavity, respectively, (p=0.054). The *H. pylori* signature was present in the saliva of subjects with negative ^13^C-UBT, RUT, and in *H. pylori* infected individuals after successful *H. pylori* eradication. We’ve also compared the prevalence of *H. pylori* both before and after *H. pylori* eradication. Pre-eradiation 7 subjects has positive saliva and 15 were negative. In 6 of the 7 with *H. pylori* pretreatment it was no longer detected by PCR after eradication. Interestingly, 9 in 15 subjects who were *H. pylori* negative before eradication had it detected post *H. pylori* eradication whereas 6 remained free of *H. pylori* after eradication.

**Figure 4:**
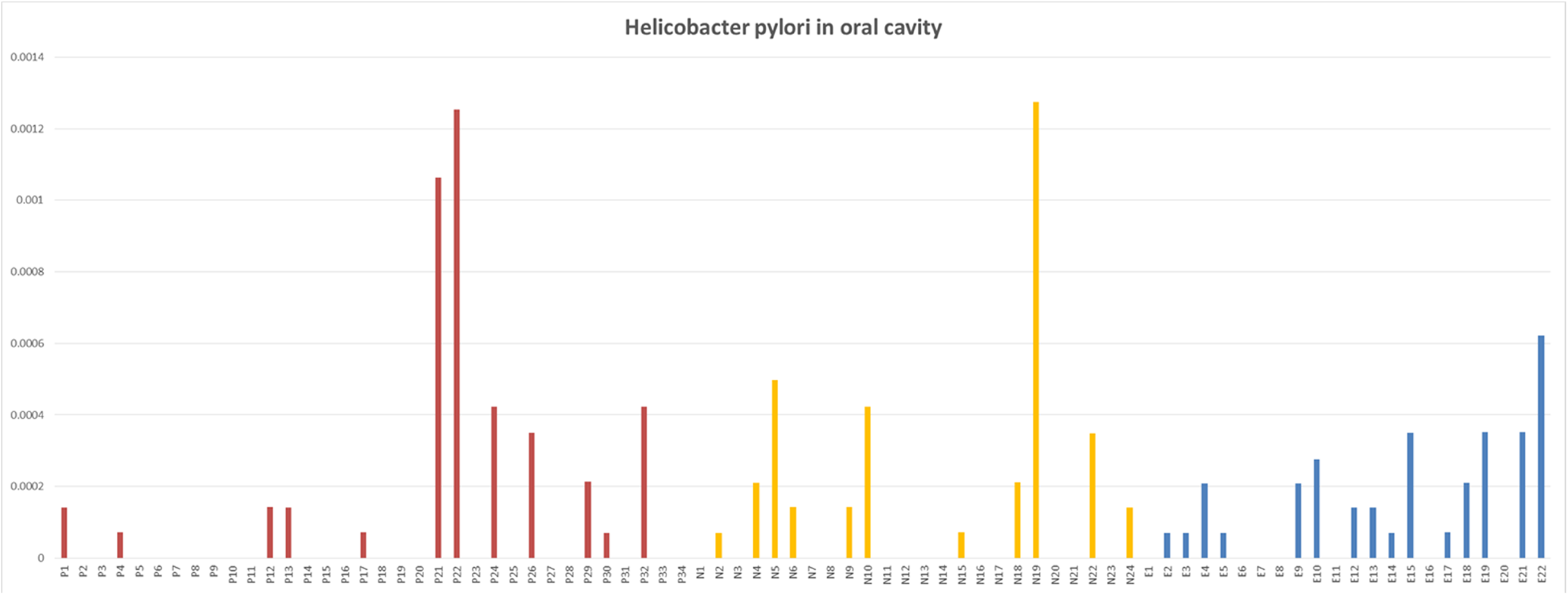
H. pylori in oral cavity of three groups. Red represent P group, yellow represent N group, and blue represent E group. N=uninfected group, P=infected group, E=eradicated group.

### Co-occurrence Network Analysis and Function Predictions

Co-occurrence analysis was used to discern relationships among the saliva microbiota at the genus level. As shown in the network diagram for the 50 most abundant genera (Figure S7), 12 genera displayed positive associations, and 1 genera displayed a negative association. Among them, *Atopobium* and *Solobacterium* exhibited a high degree of linkages with other genera. High rho values were found for the *Atopobium-Megasphaera* (0.83), *Filifactor-Treponema* (0.75), and *Atopobium-Prevotella* (0.74) pairs. Besides, *Oribacterium, Acinetobacter*, and *Ralstonia* exhibited a positive association (0.68).

To predict the functions of saliva bacterial community, PICRUSt analysis was performed based on the 16S rDNA composition data of each sample (Figure S8). A total of 41 metabolic functions were predicted in all samples with the most enrichment in membrane transport (11.8%), replication and repair (9.7%), amino acid metabolism (9.4%), carbohydrate metabolism (9.2%), translation (6.6%), and energy metabolism (5.8%).

## Discussion

A comprehensive and thorough investigation of the bacterial diversity of saliva microbiota is essential for understanding the how or whether *H. pylori* infection alters the salivary saliva microbiota. The technology of high-throughput pyrosequencing has provided new cognizance of the structures and compositions of microbiota communities.

By comparing the alpha diversity indexes we found that the bacterial diversity in saliva was similar among the *H. pylori* uninfected and *H. pylori* infected people. Our study is consistent with the notion that *H. pylori* in the stomach has little or no effect on the bacterial diversity of the oral cavity (20). The Shannon diversity index, ACE richness index, and Chao 1 richness estimator all declined after eradication of *H. pylori* compared to the baseline samples (p<0.05), which was consistent with the prior studies that use of PPIs and antibiotics may affect the oral microbiome (21, 22).

According to the beta diversity analysis based on the unweighted UniFrac distances, the community structures of saliva microbiota were different in *H. pylori* uninfected and infected individuals, which is contrary to the results of Christian’s study (20). Samples from the *H. pylori* infected subjects tended to cluster together, while the microbiota in the uninfected subjects appeared to be more variable suggesting that gastric *H. pylori* infection may affect oral bacterial components. Clear segregations by the PCoA and NMDS analysis among individuals before and after *H. pylori* eradication therapy demonstrated that successful eradication or eradication therapy changed the oral bacterial components to some extent.

In addition to the presence of different bacterial members, the abundance of some bacteria also differed significantly among groups. We clearly observed that some bacteria in the saliva of *H. pylori* infected individuals showed a significantly reduced abundance, among which *Aggregatibacter*, *Klebsiella, Fusobacterium*, and *Parvimonas* were pathogenic bacteria. *Aggregatibacter* is a dominant etiology of infective endocarditis (23). *Klebsiella* and *Fusobacterium* can lead to liver abscess, pneumonia, and meningitis (24). *Parvimonas* has been isolated as a causative agent in a variety of systemic infections, including meningitis, septic arthritis, chest wall abscess, spondylodiscitis, empyema, endocarditis, hepatic abscesses, and brain abscess (25). However, the abundance of other bacteria significantly increased in saliva of *H. pylori* infected individuals, most of which were oral microbiota composition, including *Sphingomonas*, *Ochrobactrum*, *Afipia*, *Leptotrichia*, *Oribacterium*, and *Moryell*, except *Acinetobacter* causing infectious diseases like pneumonia and urinary tract infections (26), and *Leptotrichia*, a potential cariogenic genera (27). While in Christian’s study, no significant difference in oral communities between *H. pylori* infected and uninfected individuals were detected at genus level (20), this may be due to the different target sequencing region of 16s rDNA, sample size, or geographic location. Interestingly, most *H. pylori*-enriched genera increased after the eradication, including *Ralstonia, Leptotrichia, Sphingomonas, Leptothrix, Oribacterium*, and *Acinetobacter*. The exception was *Ochrobactrum*. However, genera low expressed in *H. pylori* infected saliva experienced a further decline after *H. pylori* eradication therapy, including *Alloprevotella, Aggregatibacter*, *Leptotrichlaceae__G_1*_, *Parvimonas*, and *Fusobacterium*, most of which are pathogenic bacteria. The presence of *Ralstonia* positively correlated with the presence of *Oribacterium* and *Acinetobacter*, each of which increased in patients with *H. pylori* after successful eradication. Our study suggests that *H. pylori* infection may change the saliva microbiota by reducing the number of conditional pathogenic bacteria and increasing the number of normal bacteria composition. After *H. pylori* eradication therapy, most conditional pathogenic genera in saliva decline while most symbiotic bacteria become more abundant.

Although the clinical significance of these alterations is not known, *H. pylori* unexpectedly and clearly altered the oral microbiota composition. Previous studies have reported acid inhibition in upper gastric tract may have an effect on the oral microbiome leading to alterations in the microbiota (28). In addition, changes in gastric pH could also lead to an alteration in the pH of oral cavity (29). *H. pylori* generates large amount of urease, an enzyme which decomposes urea into ammonia and carbon dioxide and transiently reduce the acidic environment in the stomach (30). We proposed that *H. pylori* likely changed the community and structure of oral microbiota through changes in the acidic environment in stomach. The use of PPIs during the eradication therapy would further inhibit the pH in stomach, leading to further alteration in saliva microbiota, which can partially explain why genera enriched in *H. pylori* infected individuals would further increase and genera low expressed in *H. pylori* infected individuals would decline after successful eradication. Although the precise mechanism has yet to be clarified, to our knowledge this is the first study to clearly show oral microbiota alterations as a result of *H. pylori* infection in a cohort of subjects. Additional studies to investigate these possible causal relationships would likely provide interesting findings. Besides, by PICRUSt analysis, we predicted that the saliva bacterial functions mainly enriched in membrane transport, replication and repair, amino acid metabolism, carbohydrate metabolism, translation, and energy metabolism.

Using amplicon pyrosequencing of 16S rDNA V3-V4 hypervariable regions we detected *H. pylori* in the oral cavity of almost half of the subjects regardless of whether they had gastric infection with *H. pylori*. Subjects who did not have *H. pylori* in the oral cavity before eradication surprisingly had *H. pylori* detected in saliva samples after *H. pylori* eradication therapy. Clearly, using these techniques the prevalence of *H. pylori* in oral cavity is not clearly associated with colonization status in the stomach which is not consistent with the notion that the oral cavity represents a secondary site for *H. pylori* colonization (31). The gastric and oral mucosa differ markedly. For example, of the two only the gastric mucosa expresses Lewis^b^ antigen, an ABO blood group antigen, which enables adherence of *H. pylori* to the epithelial surfaces. It has been proposed that *H. pylori* is a passerby in oral cavity, rather than a colonizer and it may be also be included in the material in gastroesophageal reflux. The natural history of *H. pylori* infection has been that after *H. pylori* eradication from the stomach, gastric reinfection is rare and when it occurs early it can often be shown to be recrudescence (the same genotype) whereas later reinfections are most often reinfection with a different genotype (32). The hypothesis that *H. pylori* was a common passerby rather than a colonizer would partly explain why recurrences are most common in areas with poor sanitation and a high prevalence of *H. pylori* and rare in developed countries whose frequency of *H. pylori* infection had become much lower than that of poor regions.

## Strengths and Limitations

Our study showed that *H. pylori* infection and the eradication treatment resulted in alterations of oral microbiota. However, there were limitations to our study. The technique of high-throughput pyrosequencing we used in our study could detect the microbiota at genus level precisely. Metagenomics sequencing was not able to be used to detect *H*. pylori-specific virulence factors such as VacA, CagA, OipA, etc. or full sequence (33). One issue with the interpretation is that there was no control sample of *H. pylori* uninfected individuals receiving the same antimicrobial therapy which precluded determination about whether the presence of *H. pylori*, the antimicrobial therapy, or both were dominant factors in changing the within-individual diversity of the oral cavity.

## Conclusions

Our study showed that bacterial diversity was similar in *H. pylori* infected and uninfected people. Eradication therapy was associated with a decline the bacterial diversity in oral cavity. Both *H. pylori* infection and eradication therapy caused alterations in community and structure of the oral microbiota. *H. pylori* is found commonly in the oral cavity with no clear relation to *H. pylori* infection of the stomach.

## Acknowledgments

This study was financially supported by National Natural Science Foundation of China (81170355 and 81370592) and Clinical Research Center, Shanghai Jiao Tong University School of Medicine. We acknowledged Shanghai Personal Biotechnology Co., Ltd. For their kind help in 454 pyrosequencing and bioinformatics analysis. All authors don’t have any potentially conflicting interests.

## Supporting Information

**Table S1 Clinical parameters of the 58 subjects**. N=uninfected group, P=infected group, PE=pre-eradicated group, E=eradicated group.

**Table S2: Modified OTU table at 97% identity**. N=uninfected group, P=infected group, E=eradicated group.

**Figure S1: Length distribution of sequences determined by 454 pyrosequencing**.

**Figure S2: A classification tree showing bacterial abundance by MEGAN**. The taxonomy compositions and abundances were visualized by MEGAN (version 6.6.7). The larger the area of the colored pie chart, the greater the bacterial abundance. Different colors represent different groups, and the larger the colored sectorial area at a branch, the more the corresponding group contributed to the bacterial abundance. N=uninfected group, P=infected group, E=eradicated group.

**Figure S3: Alpha diversity (observed species number) among groups**. (A) N group and P group showed similar alpha diversity (p>0.05). N=uninfected group, P=infected group. (B) The observed species in E group were significantly lower than that of PE group and N group (p<0.01); One asterisk indicates significant differences (p < 0.05, Student’s t-test), two asterisk indicates p<0.01, three asterisk indicates p<0.001. N=uninfected group, PE=pre-eradicated group, E=eradicated group.

**Figure S4: Rank abundance curves for all OTUs**. N=uninfected group, P=infected group, E=eradicated group.

**Figure S5: Nonmetric Multidimensional Scaling (NMDS) based on unweighted UniFrac distances at the OUT level at 97% identity**. Each sample is represented by a dot. (A) The samples formed well-separated clusters corresponding to the two groups, suggesting that the bacterial structures in N group and P group were different. N=uninfected group, P=infected group. Red squares represent the N samples. Blue triangles represent the P samples. (B) Blue triangles represent the N samples. Red circles represent the E samples. The samples formed well-separated clusters corresponding to the three groups, suggesting that the bacterial structures in E group, PE group, and N group were different. N=uninfected group, PE=pre-eradicated group, E=eradicated group.

**Figure S6: Comparison of microbial variations at the genus level, using the LEfSe online tool**. (A) Histogram of the LDA scores for differentially abundant features among groups. The threshold on the logarithmic LDA score for discriminative features was set to 2.0. N=uninfected group, PE=pre-eradicated group, E=eradicated group. (B) Cladogram for taxonomic representation of significantly differences among groups. Differences are represented in the color of the most abundant taxa (red indicating N group, blue indicating PE group, green indicating E group, and white indicating non-significant). N=uninfected group, PE=pre-eradicated group, E=eradicated group.

**Figure S7: A network diagram of dominant genera showing associations among the 50 most abundant genera**. Red line represent positive associations, and green line represent negative associations.

**Figures S8: Bacterial function prediction by PICRUSt analysis**. N=uninfected group, P=infected group, E=eradicated group.

